# Global whole-genome phylogenomics of *Nakaseomyces glabratus* reveals admixture and refines sequence type-based classification

**DOI:** 10.64898/2026.04.03.716392

**Authors:** Abdul-Rahman Adamu Bukari, Brooke Sidney, Aleeza C. Gerstein

## Abstract

*Nakaseomyces glabratus* is a globally distributed opportunistic fungal pathogen. An ongoing discussion in studies of *N. glabratus* population structure has been whether genetic clusters are best defined using multilocus sequence typing (MLST) or short-read whole-genome sequencing (WGS). To assess the concordance between MLST- and WGS-based phylogenies, we analyzed a dataset of 548 *N. glabratus* WGS sequences from 12 countries. Clusters identified from WGS largely recapitulated the MLST-defined sequence type (ST) groups: fourteen WGS clusters were composed of a single MLST ST, and the remaining contained STs with very closely related MLST profiles. We thus propose a pragmatic naming convention, consistent with the system used in other microbial species, which specifies WGS cluster labels based on the primary ST. From the large WGS isolate dataset, we determined the prevalence of admixture and genomic variants. Interestingly, seven of the nine singleton isolates were admixed, in addition to 58 isolates from six different clusters. Aneuploidy was detected in 4% of isolates, most commonly in chrE, which contains ERG11, the gene encoding the enzyme targeted by azole antifungals. Aneuploid chromosomes did not exhibit elevated heterozygosity relative to the sequencing error rate, consistent with instability of extra chromosome copies. Copy number variants were found in 3% of the isolates; some of the CNVs co-occurred with aneuploidies, and were primarily identified on chrD, chrE, chrI, and chrM. Our findings demonstrate that deep splits between clusters preserve the utility of MLST ST designations for clade-level designation, yet underscore the utility of WGS for high-resolution genomic analyses.

**Article Summary:** There is an ongoing debate in studies on *Nakaseomyces glabratus* about whether traditional MLST analysis is sufficient to determine population structure, or whether the precision of whole genome sequencing (WGS) is necessary. We analyzed WGS data from 548 isolates from around the world. We found a very strong agreement between the two methods. We propose a hybrid naming system, where cluster names are based on the dominant MLST group. We used the WGS data to show that admixed isolates, and those with extra chromosomes or CNVs are rare (<7% of isolates in each class) and are distributed throughout the phylogeny.

## Introduction

*Nakaseomyces glabratus* (formerly *Candida glabrata*) is an opportunistic fungal pathogen of increasing clinical importance, particularly in the context of healthcare-associated infections among immunocompromised individuals (Gabaldón and Fairhead 2019; Li *et al*. 2021). In 2022, the World Health Organization classified *N. glabratus* as a high-priority fungal pathogen (WHO 2022). Epidemiological trends over the past two decades indicate an increasing prevalence of *N. glabratus* as commonly the second most frequent invasive yeast infection after *Candida albicans* (Pfaller *et al*. 2009, 2019; Taj-Aldeen *et al*. 2014; Chapman *et al*. 2017; Astvad *et al*. 2018; Fuller *et al*. 2019; Kord *et al*. 2020; Arastehfar *et al*. 2021). In comparison to *C. albicans*, *N. glabratus* isolates exhibit innate resistance to azoles more frequently (Oxman *et al*. 2010). *N. glabratus* infections can thus be challenging to treat, which potentially explains why they continue to increase in incidence.

*N. glabratus* belongs to the Nakaseomyces clade (Kurtzman and Robnett 2003; Fitzpatrick *et al*. 2006; Gabaldón *et al*. 2016), a group that is more closely related to the baker’s yeast *Saccharomyces cerevisiae* than to *C. albicans*. Genetic diversity among *N. glabratus* isolates has been assessed using both MLST and WGS data. MLST studies are performed by sequencing the coding regions of six conserved housekeeping genes (*FKS*, *LEU2*, *NMT1*, *TRP1*, *UGP1*, and *URA3*). These loci were chosen for strain-level typing in epidemiological and population genetic studies as they provide sufficient sequence diversity to discriminate among isolates while (Dodgson *et al*. 2005; Lott *et al*. 2010). As of May 2025, the PubMLST database (https://pubmlst.org), the comprehensive global repository for microbial molecular typing and genome diversity (Jolley *et al*. 2018), identifies 314 sequence types (STs) from 2,187 *N. glabratus* isolates.

The first study to employ WGS data examined 33 globally sampled *N. glabratus* isolates and recovered seven genetically distinct clusters, referred to as clade I through clade VII. (Carreté *et al*. 2018). The MLST and WGS topologies partially, but not completely, overlapped, demonstrating the utility of WGS in providing greater precision in identifying relatedness among isolates (Carreté *et al*. 2018). Contrary to the MLST phylogenies, which had indicated geographical enrichment in some clusters (Dodgson *et al*. 2003; Lott *et al*. 2010), the WGS phylogeny revealed no geographical structure. WGS analysis indicated a much deeper intra-clade divergence and greater between-clade genetic diversity compared to *C. albicans,* potentially suggesting that *N. glabratus* could comprise a species complex of deeply diverged lineages.

*N. glabratus* has 13 chromosomes and is typically found as a haploid. It has been widely considered asexual, as all attempts to observe mating in *N. glabratus* in the laboratory have so far been unsuccessful. However, there may be a cryptic sexual cycle, as there are distinct mating types, and mate-type switching has been observed (Gabaldón and Fairhead 2019). Most convincingly, admixture analysis using both MLST and WGS data is consistent with sexual recombination between clusters (Srikantha *et al*. 2003; Dodgson *et al*. 2005; Boisnard *et al*. 2015; Carreté *et al*. 2018; Helmstetter *et al*. 2022).

Like other fungal microbial pathogens, *N. glabratus* has a flexible genome. The first WGS study identified aneuploidies (typically disomies) in 4 of 33 isolates, predominantly in chromosomes E and G (Carreté *et al*. 2018). A larger study that used flow cytometry to analyze 500 clinical isolates found that 4 % of them were aneuploid, diploid, or polyploid (Zheng *et al*. 2022). Aneuploidies are generally thought to be lost under non-selective conditions, and are often framed as playing a transient role in fungal adaptation to drug resistance (Rustchenko 2007; Gerstein and Berman 2015). A recent experimental evolution study, however, found that fluconazole-resistant ChrE aneuploid replicates retained the aneuploidy even after 35 serial passages in YPD (i.e., drug-free medium; Ksiezopolska *et al*. 2024). This suggests that although often transient, certain aneuploidies in *N. glabratus* can be stably maintained in the absence of drug pressure, at least in the lab. Thus, the degree to which *in vitro* experiments (particularly in rich medium) recapitulate the natural environment remains unknown, however, and the stability of aneuploid chromosomes remains an open question.

To assess the relationship between phylogenetic clustering from WGS and MLST sequence types (STs), we generated new WGS and MLST phylogenies from short-read WGS data from 548 clinical isolates from 12 countries. In addition, we use the WGS data to conduct a global survey of admixture, aneuploidy, and CNVs. The resulting WGS phylogeny generally reflected ST-based groupings, yet many clusters encompassed multiple closely related STs. As MLST studies are technically less challenging and resource-intensive, and hence remain widely used, we provide a reconciliatory ground for cluster designation by naming WGS clusters by their dominant STs. We identified 65 isolates with signatures of admixture, providing additional evidence of gene flow across genetically distinct lineages, adding to the evidence of an occasional sexual cycle in *N. glabratus* in the wild. Karyotypic variation was limited. Aneuploidy and CNVs were detected in ∼4% of isolates; they were not restricted to specific phylogenetic clusters. Taken together, this study demonstrates the utility of WGS data for quantifying genomic variation, yet underscores the continued utility of MLST-based phylogenetic clustering.

## Methods

### Isolate collection

We downloaded data from 530 paired-end Illumina fastq files from 20 BioProjects from the NCBI SRA (search done on July 5, 2023); we refer to these isolates as the “*N. glabratus* WGS isolates”. Approximately half were previously published in 16 articles (Håvelsrud and Gaustad 2017; Vale-Silva *et al*. 2017; Carreté *et al*. 2018, 2019; Biswas *et al*. 2018; Barber *et al*. 2019; Guo *et al*. 2020; McTaggart *et al*. 2020; Siscar-Lewin *et al*. 2021; Xu *et al*. 2021; Szarvas *et al*. 2021; Helmstetter *et al*. 2022; Salazar *et al*. 2022; Galocha *et al*. 2022; Pais *et al*. 2022; Bukari *et al*. 2023) while 287 isolates have no publication associated with them (PRJNA596170, PRJNA593955, PRJNA329124, PRJNA524686, Table S1).

In addition, 18 isolates were acquired from the microbiology lab at Health Sciences Centre in Winnipeg, Canada in 2012 (Table S1). For those isolates, genomic DNA was extracted from single colonies using a standard phenol-chloroform protocol (Kukurudz *et al*. 2022). DNA quality and concentration were assessed spectrophotometrically (NanoDrop 2000, Thermo Scientific) and fluorometrically (Qubit 2.0 Fluorometer with the dsDNA BR Assay Kit, Invitrogen), respectively. The genomes were sequenced by the Microbial Genome Sequencing Center (Pittsburgh, USA) using the Illumina NextSeq 550 sequencing technology with paired-end reads of 150 bp. The bcl-convert v3.9.3 package (https://support-docs.illumina.com/SW/BCL_Convert/Content/SW/FrontPages/BCL_Convert.htm) was used in demultiplexing, quality control, and adapter trimming. The reads have been deposited at the National Center for Biotechnology Information (NCBI) Sequence Read Archive under BioProject ID PRJNA991137.

### *In silico* MLST typing

We assigned sequence types (STs) to the *N. glabratus* WGS isolates using stringMLST with default parameters (Gupta *et al*. 2017), leveraging its capability to retrieve the latest MLST allele and profile definitions directly from the PubMLST database (Jolley *et al*. 2018). The analysis employed the established six-locus MLST scheme for *N. glabratus* (*FKS, LEU2, NMT1, TRP1, UGP1*, and *URA3*; note that PubMLST continues to label this species as *C. glabrata*). One profile combination did not match any existing entries in the database and was submitted to the database as a new ST.

### Variant calling

The sequence reads from all WGS isolates were trimmed with Trimmomatic (v0.39) (Bolger *et al*. 2014) with standard parameters (LEADING: 10, TRAILING: 3, SLIDINGWINDOW:4:15, MINLEN: 31, TOPHRED33). Quality was assessed with FASTQC (http://www.bioinformatics.babraham.ac.uk/projects/fastqc/) and MultiQC (Ewels *et al*. 2016). Trimmed paired-end reads were mapped using bwa-mem (Li 2013) to the CBS 138 reference genome (GCA000002545v2) downloaded from the Ensembl Genome Database (Yates *et al*. 2022). The resulting SAM file was coordinate-sorted and converted to a Binary Alignment Map (BAM) file using samtools v1.9 (Li *et al*. 2009). Alignment quality was assessed with CollectAlignmentSummaryMetrics from Picard v2.26.3 (http://broadinstitute.github.io/picard) and consolidated across all samples with MultiQC (Ewels *et al*. 2016). All files had a > 95% mapping quality. BAM files were further processed with Picard to add a read group annotation so that samples with the same BioProject ID had the same read group, to remove duplicate PCR amplicons, and to fix mate pairs. The average coverage for each isolate was estimated using samtools v1.9 (Li *et al*. 2009).

Chromosomal aneuploidy and copy number variation (CNV) were quantified for each isolate. Sequence reads were aligned to the reference genome, and per-base coverage was calculated from the resulting BAM files using the samtools depth function. A custom R script (available at https://github.com/MicroStatsLab/Microstats/binCoverage.R) was then used to partition the genome into non-overlapping 5 kb bins and to calculate the average read depth per bin. The average coverage across each of the 13 chromosomes was calculated for each strain. For each isolate, the bin values were normalized to a haploid base ploidy by dividing by the median chromosome coverage.

Aneuploidies and regions with increased copy number (CNVs) were detected through visual inspection from normalized coverage profiles generated from a custom script (available at https://github.com/MicroStatsLab/Microstats/SWLine.R). When the normalized coverage deviated from an integer value for an entire chromosome or CNV region, the normalized data was re-examined assuming a diploid or triploid base ploidy (i.e., by multiplying the normalized data by 2 or 3). CNV breakpoints were manually determined as the bin where a distinct, abrupt shift in normalized read depth was observed.

The GATK Best Practices were adapted for variant calling. In sequence, HaplotypeCaller, CombineGVCFs, GenotypeVCFs, VariantFiltration, and SelectVariants (DePristo *et al*. 2011; Van der Auwera *et al*. 2013; Poplin *et al*. 2018) were used to identify single-nucleotide variants (SNPs) among all sequenced isolates in haploid mode. The resulting SNP table was hard-filtered using the suggested GATK parameters (QualByDepth < 2.0, FisherStrand > 60.0, root-mean-square mapping quality < 30.0, MappingQualityRankSumTest < −12.5, ReadPosRankSumTest < −8.0). We excluded variants that were called in known repetitive regions of the genome, as these are likely to reflect sequencing misalignments rather than true variants, i.e., the subtelomeric regions (15kb from the start and end of each chromosome), the centromeres, and the major repeat sequence regions (Table S2)

To assess heterozygosity in aneuploid chromosomes, for the 18 relevant isolates, the SNP calling analysis was repeated in diploid mode in HaplotypeCaller to determine the number of called homozygous and heterozygous SNPs after filtration. To compare the prevalence of heterozygous SNPs in aneuploid chromosomes to the number of heterozygous SNPs called due to base calling errors, we similarly re-ran 18 euploid haploid isolates, matched from the same studies, in diploid mode.

### Phylogeny construction

The multi-sample VCF file was converted to a FASTA alignment using a publicly available Python script that creates an alignment matrix for phylogenetic analysis (vcf2phylip.py v2.8, downloaded from https://github.com/edgardomortiz/vcf2phylip) (Ortiz 2019). The FASTA alignment was parsed in FastTree (2.1.11) (Price *et al*. 2010) in the double-precision mode to construct an approximate maximum-likelihood phylogenetic tree using the general time-reversible model and the -gamma option to rescale the branch lengths. The phylogeny was visualized and annotated with the Interactive Tree Of Life (iTOL, v5) (Letunic and Bork 2021).

### TreeCluster cluster designations

WGS cluster designations were determined using TreeCluster (Balaban *et al*. 2019). This tool partitions phylogenetic trees into clusters based on user-defined thresholds under eight strategies (avg clade, leaf dist max, length clade, max clade, med clade, root dist, single linkage, sum branch). Four strategies were selected for further investigation because they yielded stable cluster counts across the largest number of consecutive threshold ranges. To compare these four strategies, we selected a threshold from each range that contained the fewest singleton isolates. The clustering designations from each of the chosen strategies were then compared with the ST assignments from the *in silico* MLST analysis.

### Admixture analyses

We assessed population structure in *N. glabratus* using ADMIXTURE (Alexander *et al*. 2009) following the methods used to generate the most recent large *N. glabratus* phylogeny (n=151, Helmstetter *et al*. 2022). Input data were prepared by converting the variant call format (VCF) into PLINK BED format. Nonstandard contig names (i.e., chromosomes), which were initially labelled A-M following *N. glabratus* standard nomenclature was changed to be sequential integers (1-13) as required for ADMIXTURE. Admixture analysis was performed on haploid mode across a range of hypothetical numbers of ancestral populations (K values) from 1 to 40. Five-fold cross-validation was employed to evaluate model performance for each K; the cross-validation error outputs were parsed using a custom R script to determine the K value with the lowest error. Admixture population proportions were visualized for five K values examined in closer detail, selected based on the decline in cross-validation (CV) error, to assess ancestral contributions.

### F_ST_ calculation

Genetic differentiation between 27 *N. glabratus* WGS clusters was quantified using Hudson’s Fixation Index (F_ST_; Hudson *et al*. 1992). Hudson’s F_ST_ quantifies differentiation based on the ratio of net nucleotide divergence (i.e., differences between populations) to total nucleotide diversity (i.e., differences both within and between populations). The Hudson’s F_ST_ is robust for haploid populations and remains reliable even with low or uneven sample sizes (de Jong *et al*. 2024). The calculation was as implemented in Pixy (v1.2.7.beta1, Korunes and Samuk 2021). Pixy has been shown to better account for biases caused by treating missing genotypes. F_ST_ estimates were calculated in 10 kb windows across the genome. Mean F_ST_ values were subsequently summarized across windows to assess the level of genetic differentiation among clusters.

F_ST_ values were also calculated for each selected K using allele frequencies estimated by ADMIXTURE (Alexander *et al*. 2009), providing a measure of genetic differentiation at the inferred ancestral population level. Unlike Hudson’s F_ST_, which uses individual isolates as the unit of analysis, these estimates reflect differentiation among inferred ancestral components rather than among individual isolates. Because ADMIXTURE partitions each genome into contributions from multiple ancestral populations, this approach can reveal differentiation among ancestral components.

### Data and script availability

The sequenced files have been deposited on SRA with the BioProject ID PRJNA1337144. All data, supplementary tables, and scripts to run statistical analyses and generate figures are available at https://github.com/microstatslab/Nglabratus_phylogenetics.

## Results

To determine how well MLST-based classifications recapitulate WGS phylogenetic relationships, we analyzed WGS and MLST data from 548 *N. glabratus* isolates. The data set is reflective of geographic bias in the data that has been submitted to NCBI; although isolates were collected from four continents, the majority came from North America (59.0%) and Europe (31.4%), followed by Oceania (9.1%), and Asia (0.4%) (Figure 1). Similarly, although the isolates were obtained from a range of clinical sources, over three-quarters came from the circulatory system (79.2%). Additional sources included the gastrointestinal tract (2.6%), oral cavity (2.4%), and urogenital tract (urine, kidney, urinary tract; 1.8%), with less than 1% of isolates from other sites (Table S1; note that the source was missing for 53 isolates).

**Figure 1.**
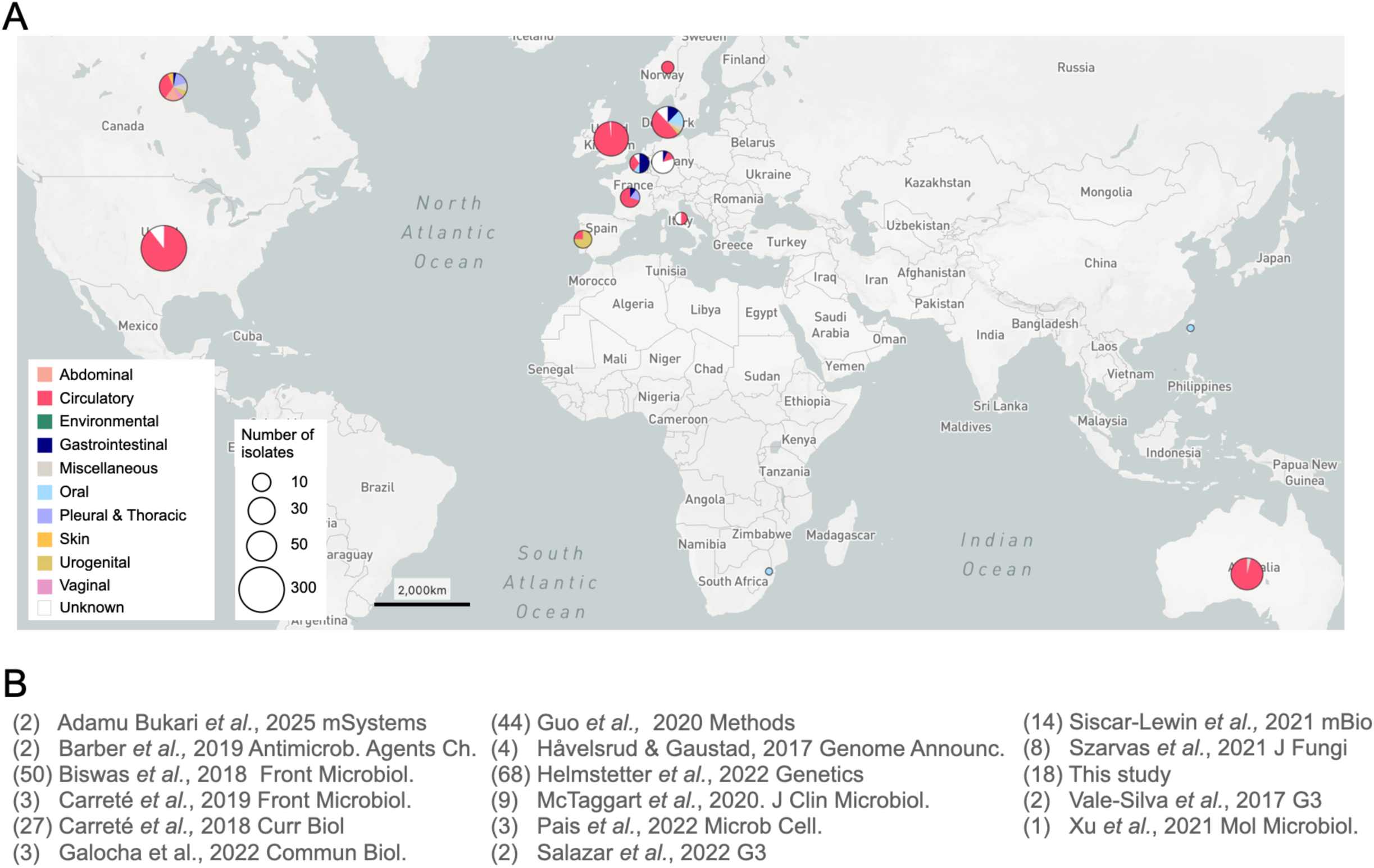
Distribution and literature source of isolates used in this study. A) Geographical distribution of the 548 isolates analyzed in this study. The circle sizes indicate the log of the number of isolates in each region, while colored sectors represent the respective isolation sources. The image was generated using Microreact. B) The publication source of isolates, the number of isolates from each manuscript is provided in brackets. 287 isolates were not linked to publication.

### *N. glabratus* phylogenetic clusters

A total of 82,564 SNP positions were identified across all isolates and used to construct the phylogeny. To identify clusters, we applied eight tree-based clustering strategies in TreeCluster (Balaban *et al*. 2019), each across the range of possible threshold values. The number and composition of potential clusters varied by method and threshold space (Figure 2A), with the number of identified clusters ranging from 1 to 200. Three different strategies identified 27 clusters, each across many consecutive threshold values (single linkage, 379 thresholds; maximum clade, 353 thresholds; length clade, 219 thresholds; Figure 2A). The next best supported number of clusters was 18, identified from the medium clade strategy across 375 consecutive thresholds (Figure 2A). Visually, the nine clusters that were lost were distinct MLST clusters that were merged with other clusters in the medium clade strategy (Figure 2B).

**Figure 2:**
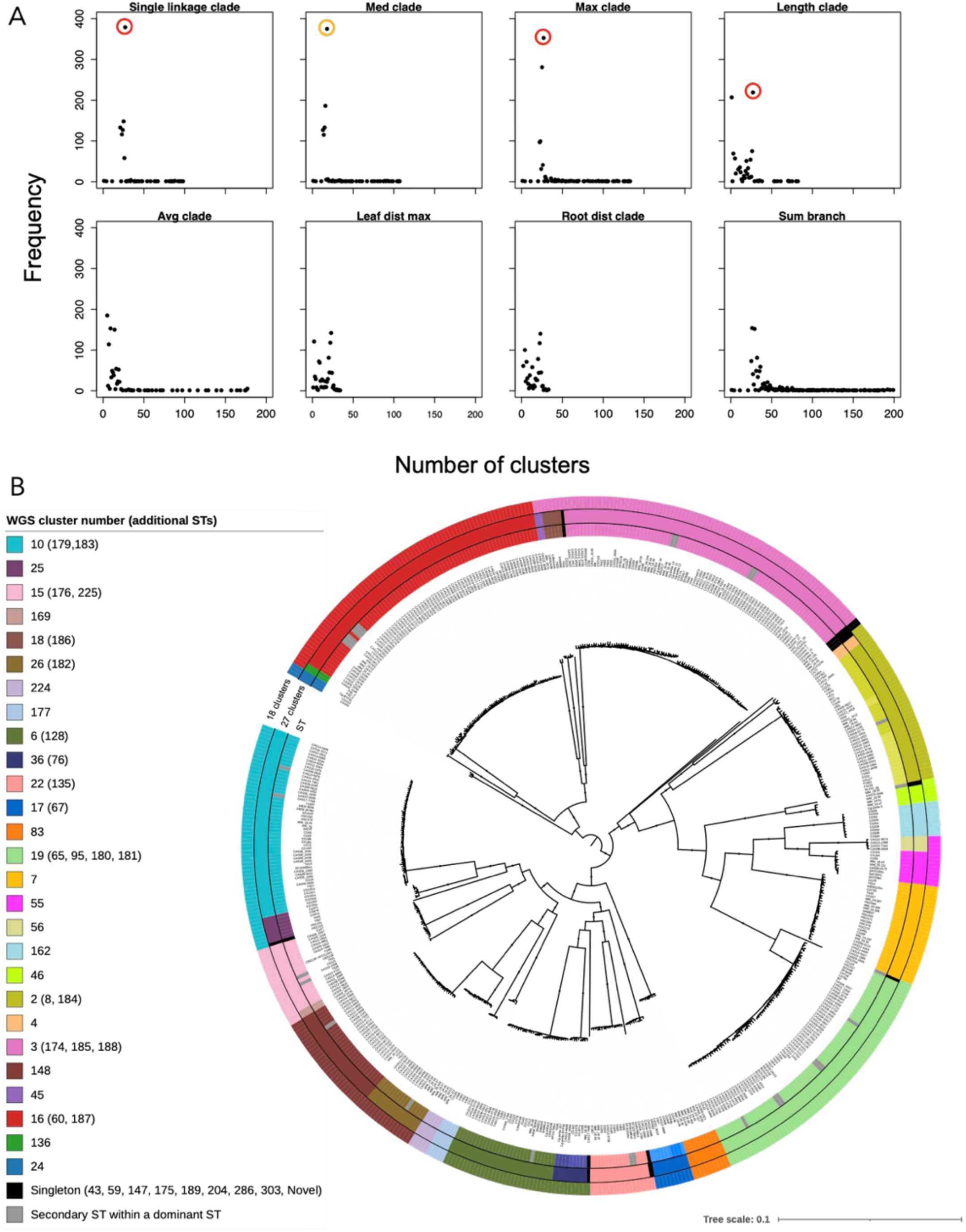
Clustering consistency and phylogenetic structure of 548 *N. glabratus* isolates. (A) Cluster prediction results using TreeCluster across eight different strategies, showing the number of clusters identified across 1000 distance thresholds (thresholds incremented in units of 0.001). Three strategies (single linkage clade, max clade and length clade) identified 27 clusters across many thresholds (red circles). The med clade strategy also showed considerable threshold stability with 18 clusters (orange circle). (B) Maximum likelihood phylogeny of the 548 isolates. The inner ring shows sequence type (ST) assignments from the pubMLST database. The middle and outer rings display the 27 and 18 cluster designations, respectively. In clusters containing multiple STs, the number listed in the legend is the predominant ST (which we propose using as the WGS cluster name). When multiple ST numbers are listed on the same row, they cluster together in the WGS phylogeny with the proposed WGS cluster number, i.e., the predominant ST group, listed first. Black bubbles on branches represent bootstrap support ≥ 0.9. The phylogeny was rooted at the midpoint

The majority of threshold space consistent with 27 clusters identified nine singleton isolates (single linkage: 0.004-0.0354; max clade: 0.099-0.0361; length clade: 0.023-0.018). A smaller amount of threshold space identified seven or eight singletons. To evaluate the differences among potential phylogenies, we visualized phylogenetic assignments for the minimum and maximum threshold value identifying seven, eight, or nine singletons from each of the three strategies. The same two isolates (HSC003 and WM_18.50) were repeatedly implicated as either being classified as singletons or grouped into clusters, depending on the strategy and threshold (Figure S1). These isolates appear visually distinct on the phylogeny. Accordingly, we proceeded with the majority-supported, 27-cluster, 9-singleton isolate tree (Figure 2B, middle ring).

The WGS isolates fell into 57 STs, and the sequence type distribution closely matched the 27 cluster-isolate designations from TreeCluster (Figure 2B). 15 of the 27 clusters contained isolates from a single ST group. 11 of the 14 remaining clusters contained isolates from one dominant ST group alongside single or a small number of isolates from secondary STs; in each case, isolates from the secondary STs differed from the predominant ST MLST scheme in only one of the six genes (Table S3). Most of the MLST differences between isolates in dominant and secondary ST groups that clustered closely together occurred at the *NMT1* (14/31) and *LEU2* (8/31) loci. The other four genes were variable in fewer than five isolates (Table S3). Three clusters (ST2/ST8, ST17/ST67 and ST36/ST76) contained two dominant STs that contributed approximately equal numbers of isolates; each differed in only one gene (UGP1, TRP1 and LEU2, respectively). The nine singleton isolates from the WGS phylogeny all had unique STs. From this point onward, the 27 clusters are each referred to by their dominant ST group. For the three clusters with two dominant groups, we compared the number of isolates assigned to each ST group in pubMLST and used the number with the most isolates; in each case, this was the lower ST number. Although sampling bias cannot be ruled out, several clusters were composed entirely of isolates from a single place (e.g., cluster 136 isolates were exclusively from Asia) or a single region, suggesting a strong geographic association for specific STs (Table 1).

**Table 1:**
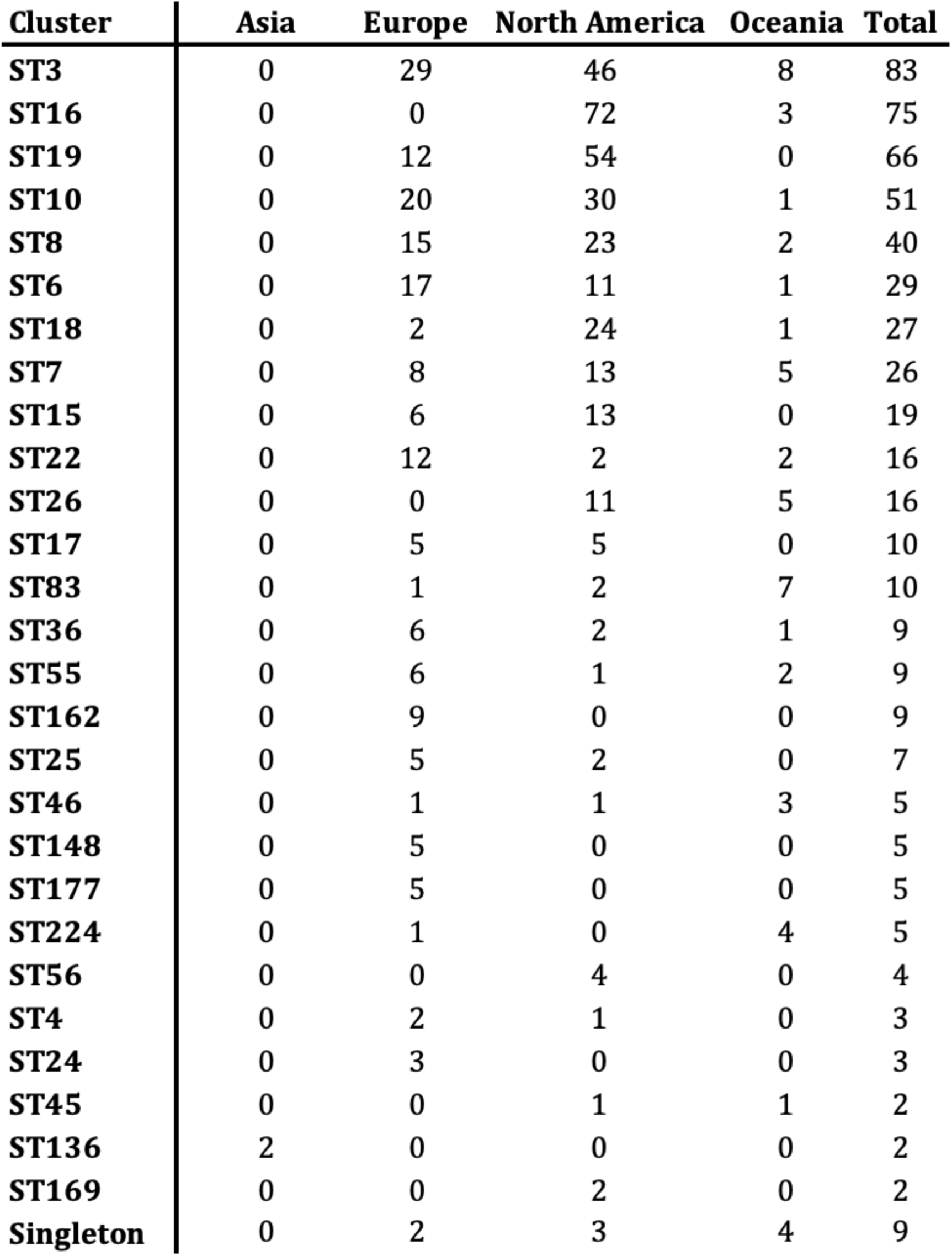
Distribution of *N. glabratus* clusters across continents.

| Cluster | Asia | Europe | North America | Oceania | Total |
| --- | --- | --- | --- | --- | --- |
| ST3 | 0 | 29 | 46 | 8 | 83 |
| ST16 | 0 | 0 | 72 | 3 | 75 |
| ST19 | 0 | 12 | 54 | 0 | 66 |
| ST10 | 0 | 20 | 30 | 1 | 51 |
| ST8 | 0 | 15 | 23 | 2 | 40 |
| ST6 | 0 | 17 | 11 | 1 | 29 |
| ST18 | 0 | 2 | 24 | 1 | 27 |
| ST7 | 0 | 8 | 13 | 5 | 26 |
| ST15 | 0 | 6 | 13 | 0 | 19 |
| ST22 | 0 | 12 | 2 | 2 | 16 |
| ST26 | 0 | 0 | 11 | 5 | 16 |
| ST17 | 0 | 5 | 5 | 0 | 10 |
| ST83 | 0 | 1 | 2 | 7 | 10 |
| ST36 | 0 | 6 | 2 | 1 | 9 |
| ST55 | 0 | 6 | 1 | 2 | 9 |
| ST162 | 0 | 9 | 0 | 0 | 9 |
| ST25 | 0 | 5 | 2 | 0 | 7 |
| ST46 | 0 | 1 | 1 | 3 | 5 |
| ST148 | 0 | 5 | 0 | 0 | 5 |
| ST177 | 0 | 5 | 0 | 0 | 5 |
| ST224 | 0 | 1 | 0 | 4 | 5 |
| ST56 | 0 | 0 | 4 | 0 | 4 |
| ST4 | 0 | 2 | 1 | 0 | 3 |
| ST24 | 0 | 3 | 0 | 0 | 3 |
| ST45 | 0 | 0 | 1 | 1 | 2 |
| ST136 | 2 | 0 | 0 | 0 | 2 |
| ST169 | 0 | 0 | 2 | 0 | 2 |
| Singleton | 0 | 2 | 3 | 4 | 9 |

The 27 clusters are deeply branching from one another, explaining why MLST analysis does such a good job of recapitulating the WGS phylogeny for *N. glabratus*. Genetic distance between isolates from the two most distant clusters (clusters 3 and 56: average 21 SNPs/kb between isolate pairs) is an order of magnitude higher than that between the most closely related ones (clusters 18 and 26 and clusters 55 and 56: average of 4 SNPs/kb). The genetic distance between closely related clusters is still two orders of magnitude larger than the genetic divergence within clusters (0.04 - 1.11 SNPs/kb, Table S4). Comparatively, the level of variation between closely-related *N. glabratus* clusters is higher than the degree of genetic variation among distant clusters in *C. albicans* (average of 3.7 SNPs/kb) (Hirakawa *et al*. 2015). Consistent with these patterns, pairwise F_ST_ values among the clusters ranged from 0.49 (cluster 18 vs. cluster 26) to 0.99 (cluster 56 vs. cluster 136), indicating extremely high genetic differentiation (Figure S2A). Generally, the pairwise Hudson’s F_ST_ values among the *N. glabratus* clusters were strongly skewed toward high differentiation. The mean F_ST_ was 0.96 and the median 0.97, indicating that most cluster pairs are highly diverged (Figure S2B).

### Admixture analyses

We identified admixed isolates with an unsupervised model-based clustering using ADMIXTURE. As there was no clear K to use (Figure 3A), we first examined K = 27, as this matched the number of WGS clusters (Figure 3A). To assess the number of ancestral populations contributing to each isolate, we counted all ancestry components contributing 1% or more of the genome (i.e., q ≥ 0.01). The majority of isolates (483 of 548) were classified as single-ancestry. We thus identified 65 admixed isolates, representing ∼12% of all isolates (Figure 3B, Table S5). The majority (50) had contributions from two ancestral populations, while three isolates had ancestry from three populations, two from four, and one from five. The remaining nine isolates had contributions from seven or more populations. Across all admixed populations, fourteen isolates had a dominant ancestry component contributing more than 75% of their genome, while 32 isolates had the major component contributing between 50% and 75%. Thus, the vast majority of isolates derive their ancestry predominantly from a single population, while a small subset exhibits more complex patterns of admixture involving multiple ancestral sources. Evidence of admixture was not equally detected among clusters. All of the isolates within 18 clusters shared a single ancestor (Figure 3C). The majority of admixed isolates were from cluster 3 (32/83 isolates; three ancestries) and cluster 19 (14/67 isolates; two ancestries), with small numbers from cluster 16 (6/75 isolates; two ancestries), cluster 136 (2/2 isolates; two ancestries), cluster 169 (2/2 isolates; over ten ancestries), and cluster 45 (2/2 isolates; four or five ancestries). All singletons had highly admixed ancestries. Isolates in clusters 55 and 56 originate from the same ancestral population but differ at two loci (*LEU2* and *URA3*), indicating minor divergence. Overall, these patterns reveal a spectrum of population structure, ranging from genetically homogeneous lineages to highly admixed groups, underscoring the complex evolutionary history of *N. glabratus*.

**Figure 3:**
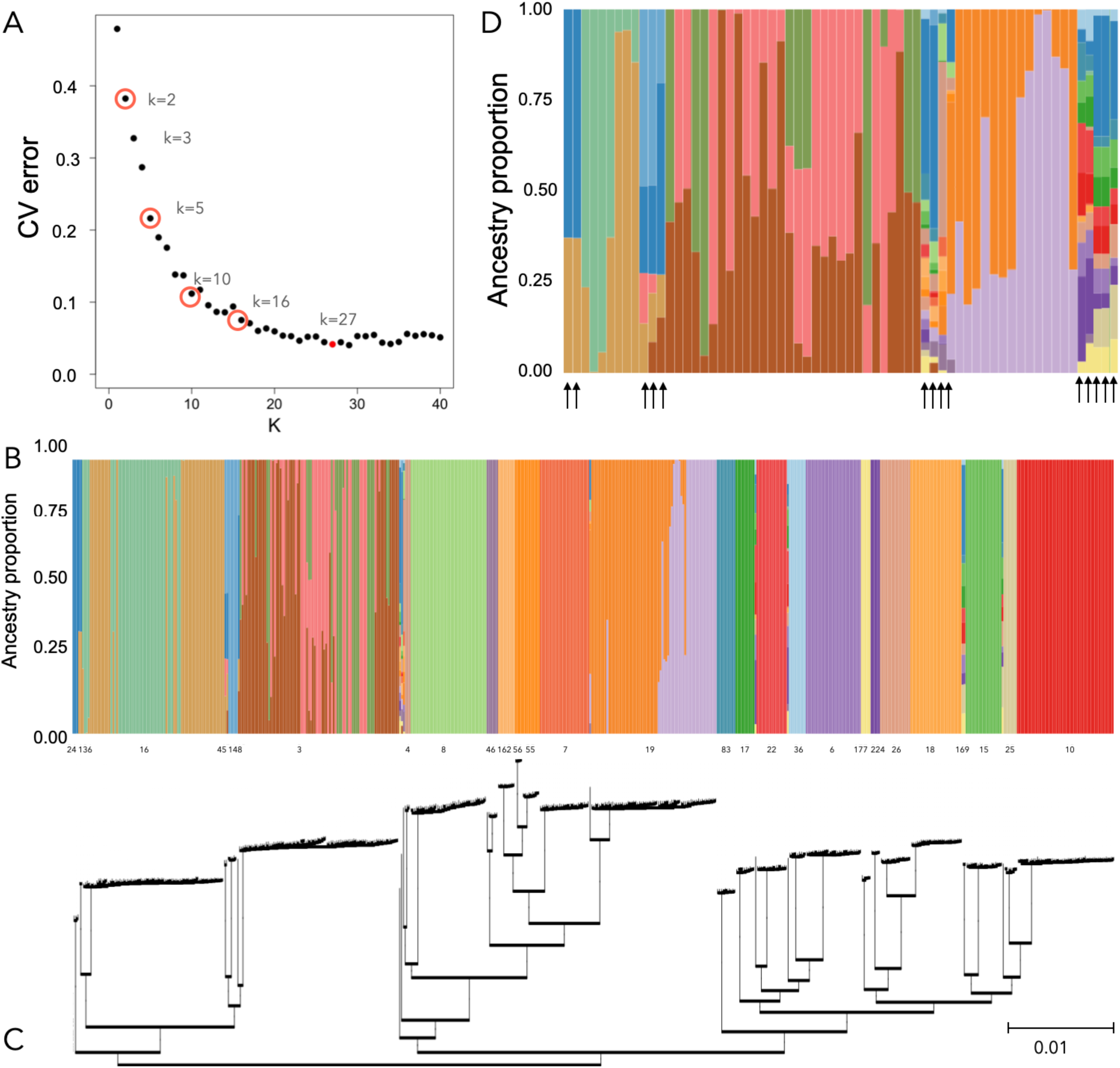
Population structure of *N. glabratus* inferred from genome-wide SNP data. Population structure of N. glabratus inferred from genome-wide SNP data. A) Cross-validation (CV) error from unsupervised ADMIXTURE analysis of variant sites across the *N. glabratus* population, testing K-values from 1 to 40. We compared admixture plots from the circled k-values, focusing on k =27, which had the lowest CV error and matched the number of MLST clusters. B) ADMIXTURE plot of all isolates at k = 27, revealing 65 individuals with evidence of mixed ancestry. (C) The maximum likelihood phylogeny constructed with FastTree was used to order the isolates. (D) A closer look at the 65 admixed isolates with arrows indicating the 14 isolates which were consistently admixed in all admixture analysis k-values.

The regional variation in admixed isolates generally followed the sampling bias, with the majority originating from North America (31 isolates), followed by Europe (17), Oceania (5), and Asia (2). Admixed isolates were recovered from diverse clinical sources, again following sampling bias; most are from the circulatory system (33 isolates), with other isolation sites including unknown sources (13), abdominal samples (3), oral sites (3), urogenital tract (2), and the gastrointestinal tract (1 isolate). This distribution suggests that admixed lineages are not clinically restricted to specific body sites.

In addition to K = 27, we analyzed the ancestries of isolates at four other K-values (2,5,10,16), selected based on the largest drops in the cross-validation scree plot (Figure 3A; Figure S5). Of the 65 isolates from K =27 that were identified as admixed, 14 were consistently identified to be of mixed ancestry across all examined K-values, eight were admixed in one additional K-value, while 43 were not identified as admixed in the others. The consistently admixed isolates are from multiple clusters (6/14) or are singletons (8/14). Pairwise F_ST_ among inferred ancestral populations were calculated for each selected K to assess genetic differentiation following admixture analysis. Mean FST values increased with higher K. From K = 6 onward, mean F_ST_ values exceeded 0.7, reaching 0.91 at K = 27, consistent with pronounced divergence among inferred populations (Figure S2C).

### Karyotypic variation

Aneuploidy was detected in 22 isolates (4%). Disomies were observed in six chromosomes: chrE in 14 isolates, chrA in 3 isolates, chrD in 3 isolates, chrC in 2 isolates, chrF in one isolate and chrL in one isolate. Most aneuploid isolates exhibited a single gain, yet three isolates had multiple aneuploidies (one with disomies in chrC and chrE, one with disomies in chrA and chrD, and one with disomies in chrA, chrC, and chrE). Karyotypic variation was not restricted to a single cluster (Figure S4). To genomically assess aneuploid chromosome stability, we compared heterozygosity levels between aneuploid and non-aneuploid chromosomes. Aneuploid chromosomes all exhibited a similar number of predicted heterozygous SNPs compared to euploid haploid chromosomes, where heterozygous base calls reflect a variant-calling error (Figure 4). This indicates that the aneuploid chromosomes in this *N. glabratus* WGS isolate set are recent, i.e., substantial *de novo* heterozygous variation has not accumulated. Copy number variation (CNV) was also detected in 14 isolates, including five with aneuploidy in different chromosomes (i.e., in those five isolates, the CNVs did not occur in the aneuploid chromosomes; Figure S3). The CNVs ranged from 30 to 810 kb and were found on chromosomes D, E, I, K, L and M.

**Figure 4:**
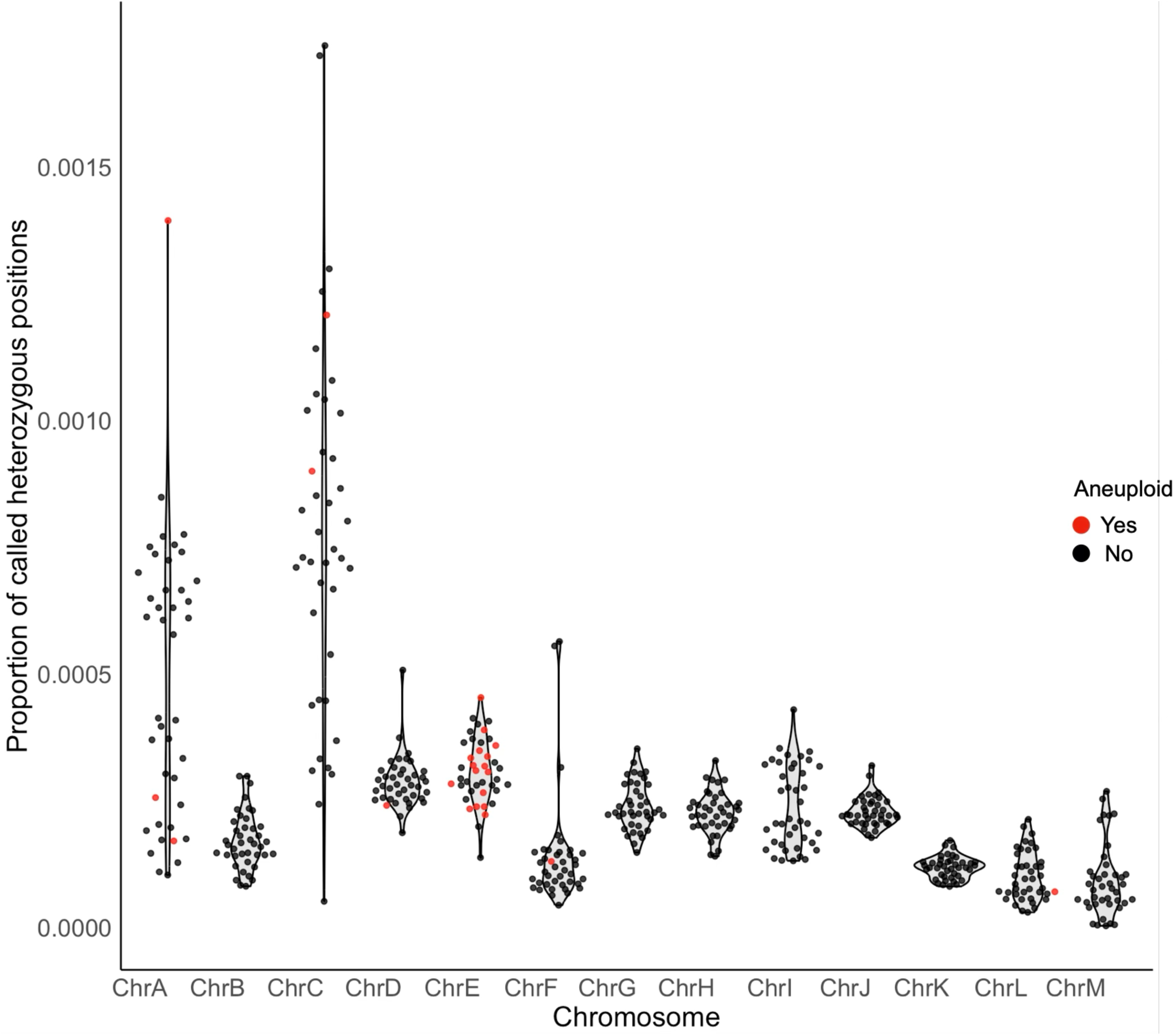
Distribution of homozygous and heterozygous SNPs in aneuploid isolates called in diploid mode. Levels of heterozygosity per chromosome are similar for both aneuploid and non-aneuploid isolates.

## Discussion

In this study, we sought to provide a harmonized approach to cluster designation in *N. glabratus.* Our analysis of 548 isolates revealed that WGS-defined clusters correspond closely with MLST-defined topology. Fourteen WGS clusters consist exclusively of a single ST, while mixed clusters typically include a dominant ST and one or a few closely related STs with few variants differentiating them. Given these close genetic relationships, we propose that clusters containing multiple STs be labelled according to the dominant ST. This approach aligns with pragmatic practices in microbial genomics, such as in *Listeria monocytogenes*, where clonal complexes are defined as groups of STs differing by one allele, and the “complex” takes the name of the largest or first-described ST (Ragon *et al*. 2008; Chenal-Francisque *et al*. 2011; Haase *et al*. 2014). A similar designation system is also used in *Staphylococcus aureus* (Feil *et al*. 2003), where clonal complexes are defined as groups of STs that match a central genotype at four or more loci, unless they are more closely related to a different central genotype (https://pubmlst.org/organisms/staphylococcus-aureus/clonal-complexes). Such naming promotes clarity and facilitates communication across studies.

This approach is different from what has previously been done in *N. glabratus* WGS studies. Dodgson et al. (Dodgson *et al*. 2003) and Carreté et al. (Carreté *et al*. 2018) both advocated for the use of Roman-numeral clade naming, while Helmstetter et al. (Helmstetter *et al*. 2022) supported ST-based nomenclature tied strictly to MLST profiles. We believe that the naming strategy proposed here strikes a balanced middle ground. By retaining cluster names based on the primary ST’s label, we reconcile precision with practicality. Within clusters containing multiple STs, deviation from the primary ST label was mainly due to variability in the NMT1 locus. This aligns with previous observations highlighting NMT1 as the most diverse among the six loci used in the MLST scheme (Dodgson *et al*. 2003; Gabaldón *et al*. 2020).

The magnitude of genetic differentiation observed among the 27 *N. glabratus* clusters (pairwise Hudson’s F_ST_ = 0.49–0.99) indicates pronounced population structuring within this species. This closely aligns with previous genome-wide F_ST_ estimates reported among *N. glabratus* sequence types (0.64–0.90; Helmstetter et al. 2022). According to Wright’s qualitative guidelines, values exceeding 0.25 represent “very great genetic differentiation,” and all cluster pairs surpass this threshold (Hartl and Clark 2007). The exceptionally high F_ST_ values observed in *N. glabratus* exceed (or approach the upper bound of) differentiation reported in many other fungal pathogens. For example, comparisons between major *Candida albicans* genetic clusters (clusters 1, 2, 3, 4, 11) show a pairwise F_ST_ range of 0.79–0.83. When these five large *Candida albicans* clusters are compared to clade 13, which is likely a cryptic species (*C. africana*, Romeo and Criseo 2011), F_ST_ values ranged from 0.87-0.90 (Ropars *et al*. 2018). F_ST_ values associated with cryptic species boundaries in other fungi overlap with the values we found. In *Paracoccidioides brasiliensis*, cryptic species exhibit pairwise F_ST_ values of 0.63–0.89 (Matute *et al*. 2006). The similarity between these cryptic-species-level divergences and the upper range of F_ST_ within *N. glabratus* reinforces the interpretation that this species complex harbors multiple deeply diverged, evolutionarily independent lineages. Although we do not interpret these 27 clusters as corresponding to 27 distinct species, the exceptionally high divergence among several cluster pairs suggests the presence of deeply separated evolutionary lineages. An attempt to include an outgroup (*C. bracarensis* isolate CBM5ILL) was not informative, as it clustered within ST16 and was therefore excluded from the phylogeny. Whether new species should be designated from isolates currently classified as *N. glabratus* requires further community discussion and evaluation.

We identified 65 of 548 isolates with evidence of admixture when we set 27 ancestral groups. Of these, 14 were supported from analysis with four different ancestral groups (2, 5, 10, 16). Most admixed isolates were within cluster 3, a cluster previously associated with admixture (Carreté *et al*. 2018), while 13 admixed isolates were found in cluster 19, which had not been previously identified as mixed-ancestry. In analyzing 181 isolates, Helmstetter et al. (2022) identified five admixed isolates from K=20 (2.8 %), three of which were confirmed in this study (CG57, WM_18.66, WM_05.155). Of the two isolates remaining, one (CG1), was not detected as admixed in our analyses. The other isolate, M17, was not available for download at the time of the analyses. Admixed isolates were geographically widespread, which may serve to maintain or enhance genetic diversity within *N. glabratus* (Steensels *et al*. 2021). A peculiar observation in this study as well as other was the observation that the admixture K used as optimal was closely similar to the number of clusters observed in the phylogeny (Carreté *et al*. 2018; Helmstetter *et al*. 2022). This is most likely because of the violation of loci independence, which is contrary to the assumptions underlying admixture analyses. Linkage disequilibrium analysis based on the six MLST loci showed a positive standardized index of association (0.2492) and a variance of pairwise differences exceeding that expected under linkage equilibrium (2.0530 vs. 0.9989), indicating a clonal population structure and the presence of linkage disequilibrium in *N. glabratus* (Xu *et al*. 2025). We are thus very cautious in over-interpreting the results. The rarity and small size of these admixed groups indicate minimal introgression and restricted gene flow among major clusters, further supporting the hypothesis that the current *N. glabratus* isolate collection could represent a species complex.

Our findings highlight the influence of geographic sampling bias on the observed distribution of *N. glabratus* STs. Our identification of ST16 as overrepresented in North America aligns with others (Kuthan 2025), as does the broad distribution of ST3 across multiple continents. However, while Dodgson et al. (2003) reported an enrichment of ST19 in Europe, our study and that of Biswas et al. (Biswas *et al*. 2018) found it to be more prevalent in North America. Similarly, the absence of Asian ST7 isolates in our study (previously described as enriched in Japan, Dodgson *et al*. 2003) likely reflects a lack of WGS data from that region. All current datasets are skewed toward North America and Europe, limiting our ability to draw firm conclusions about global primary ST distribution. A concerted effort to include underrepresented regions will be essential to resolve conflicting findings and to better understand the population structure of *N. glabratus* populations.

Aneuploidy was detected in 22 (4 %) isolates, consistent with others (Zheng *et al*. 2022). ChrE carries the *ERG11* gene (the target of azole drugs) and its amplification has been shown to correlate with resistance phenotypes (Marichal *et al*. 1997). This is similar to *C. albicans* where amplification of chromosome 5 (which harbours the *ERG11* and *TAC1* loci) confers a reversible azole resistance (Selmecki *et al*. 2009; Vande Zande *et al*. 2023). Our analyses also show that aneuploid chromosomes lack elevated heterozygosity, which supports their classification as recent *de novo* events (Carreté *et al*. 2018). This pattern is mirrored in other studies where aneuploidy emerges under stress and reverts during relaxed conditions. Furthermore, the presence of CNVs (both segmental deletions and amplifications) spanning multiple chromosomes, with co-occurrence alongside aneuploidies in 5 of 14 CNV-containing isolates (36%), denotes an additional layer of genomic plasticity.

## Conclusion

Taken together, our findings demonstrate that while MLST remains broadly concordant with WGS-based phylogenetic structure, whole-genome data can detect and resolve fine-scale sequence type nuances within *N. glabratus* phylogenies. The proposed ST-based cluster naming provides a pragmatic alternative to previous inconsistent nomenclatures and is supported by practices in other microbial systems. The deep phylogenetic divergence observed, together with variations in aneuploidies and CNVs highlights the genomic diversity of *N. glabratus*. The high F_ST_ values among clusters, approaching or exceeding those observed between cryptic fungal species and limited admixture, suggest long-term reproductive isolation and limited gene flow, indicating that *N. glabratus* might be a species complex.

## Acknowledgements

We thank Shared Health, Diagnostic Services Manitoba, and Dr. Markus Stein for facilitating the acquisition of clinical strains from the Health Sciences Centre in Winnipeg, Manitoba. We are grateful to the members of the MicroStats lab over the years for their valuable comments and feedback throughout this project. Computational analyses were supported in large part by Ali Kerrache (Prairies Region) and by resources provided through the Digital Research Alliance of Canada (https://www.alliancecan.ca/). This work was funded by the Natural Sciences and Engineering Research Council of Canada (RGPIN-2019-05867). A.C.G. acknowledges support from the CIFAR Azrieli Global Scholars Program and start-up funding from the University of Manitoba. A.-R.A.B. was supported by an EvoFunPath (NSERC CREATE) fellowship.

## Supplementary information

**Figure S1.**
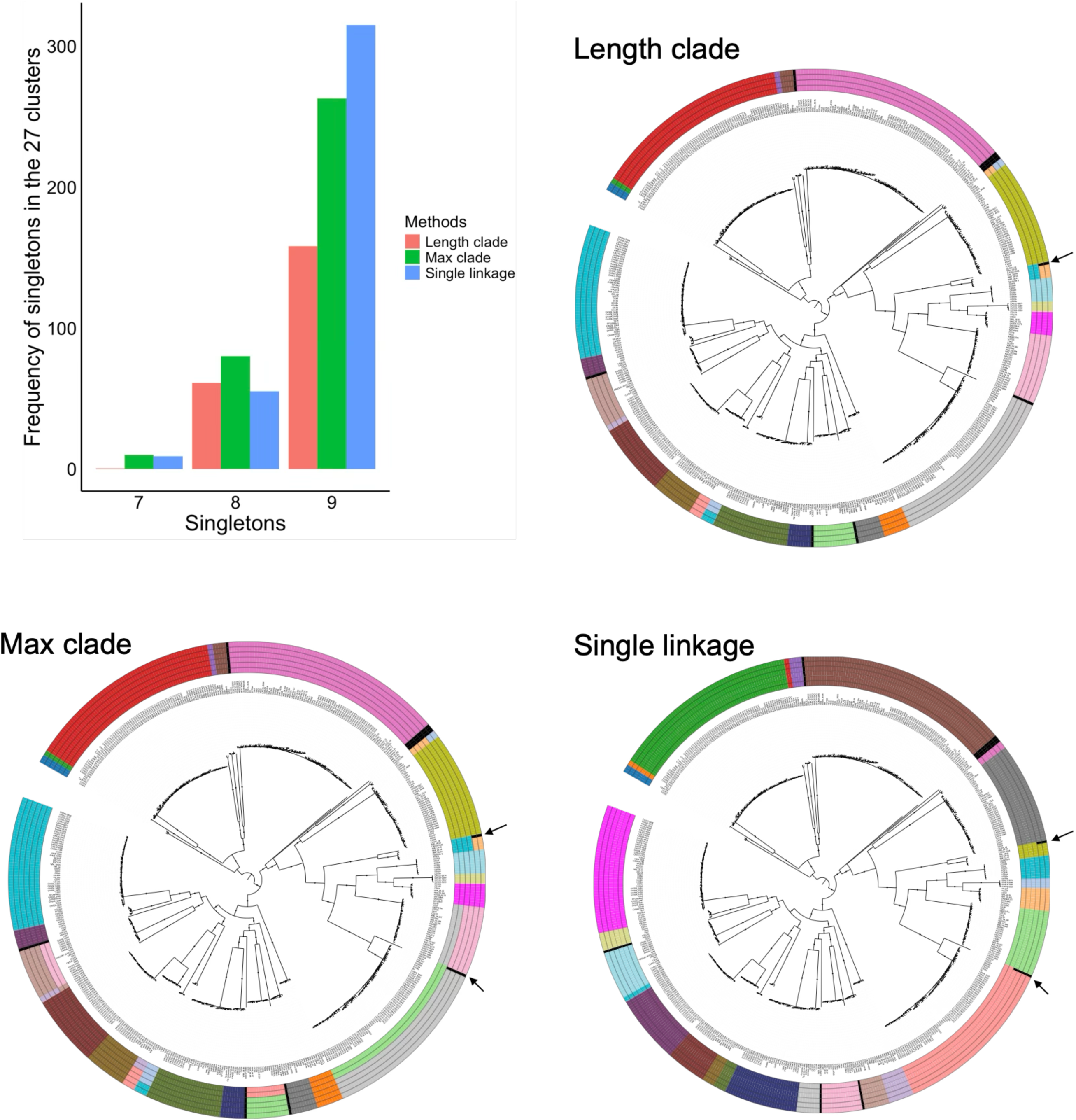
Comparison of singleton distributions within the 27 clusters predicted by three TreeCluster strategies across consecutive branch length thresholds (length clade: 0.0023–0.018; max clade: 0.0099–0.0361; single linkage: 0.00354–0.004) across the *N. glabratus* phylogeny. (A) Frequency of singletons in the predicted 27 clusters across the three TreeCluster methods. (B–D) Phylogenetic positions of isolates that contributed to variation in singleton counts under each clustering strategy. For each strategy, cluster designations corresponding to threshold boundaries with shared singleton counts are shown as colored concentric rings around the phylogeny. Length clade has four such boundaries (due to having only two singleton groups, 8 and 9, within the 27 predicted clusters), whereas max clade and single linkage each have six boundaries (due to having three singleton groups, 7, 8, and 9, within the 27 predicted clusters). Arrows indicate the two isolates responsible for deviation from the dominant 9-singleton frequency.

**Figure S2.**
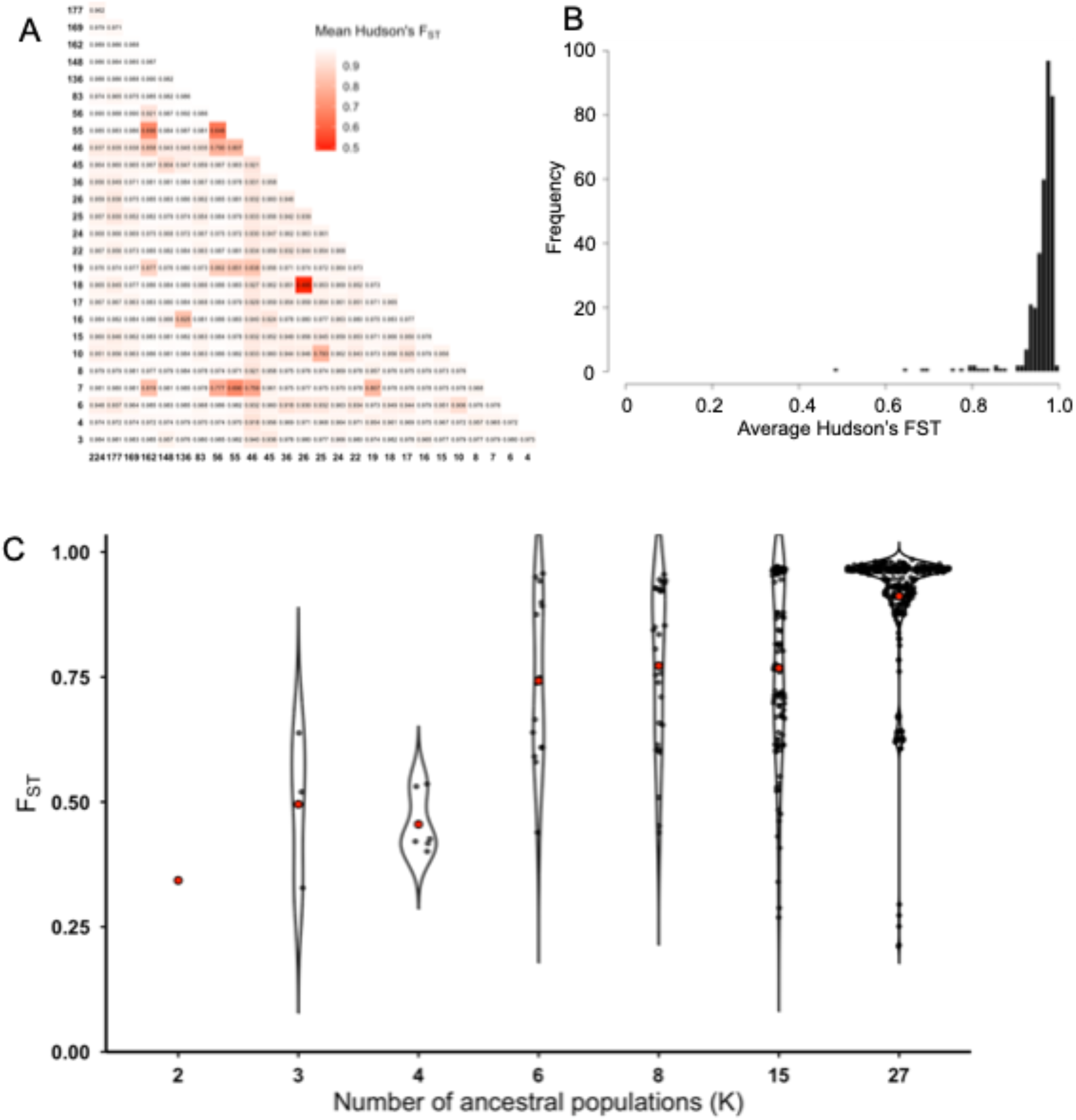
Fixatation indices of the clades and ancestral populations. A) The Mean Hudson’s FST between the *N. glabratus* clades. Pairwise FST values indicate high genetic differentiation, ranging from 0.485 to 0.99. Heatmap colors range from red (low FST) to white (high FST), representing the degree of genetic divergence between clades. B) Distribution of Hudson’s FSTs in comparing the various *N. glabratus* clusters. C) The distribution of pairwise FSTs between ancestral populations within each K. Each point represent an comparison and the red points represents the mean.

**Figure S3.**
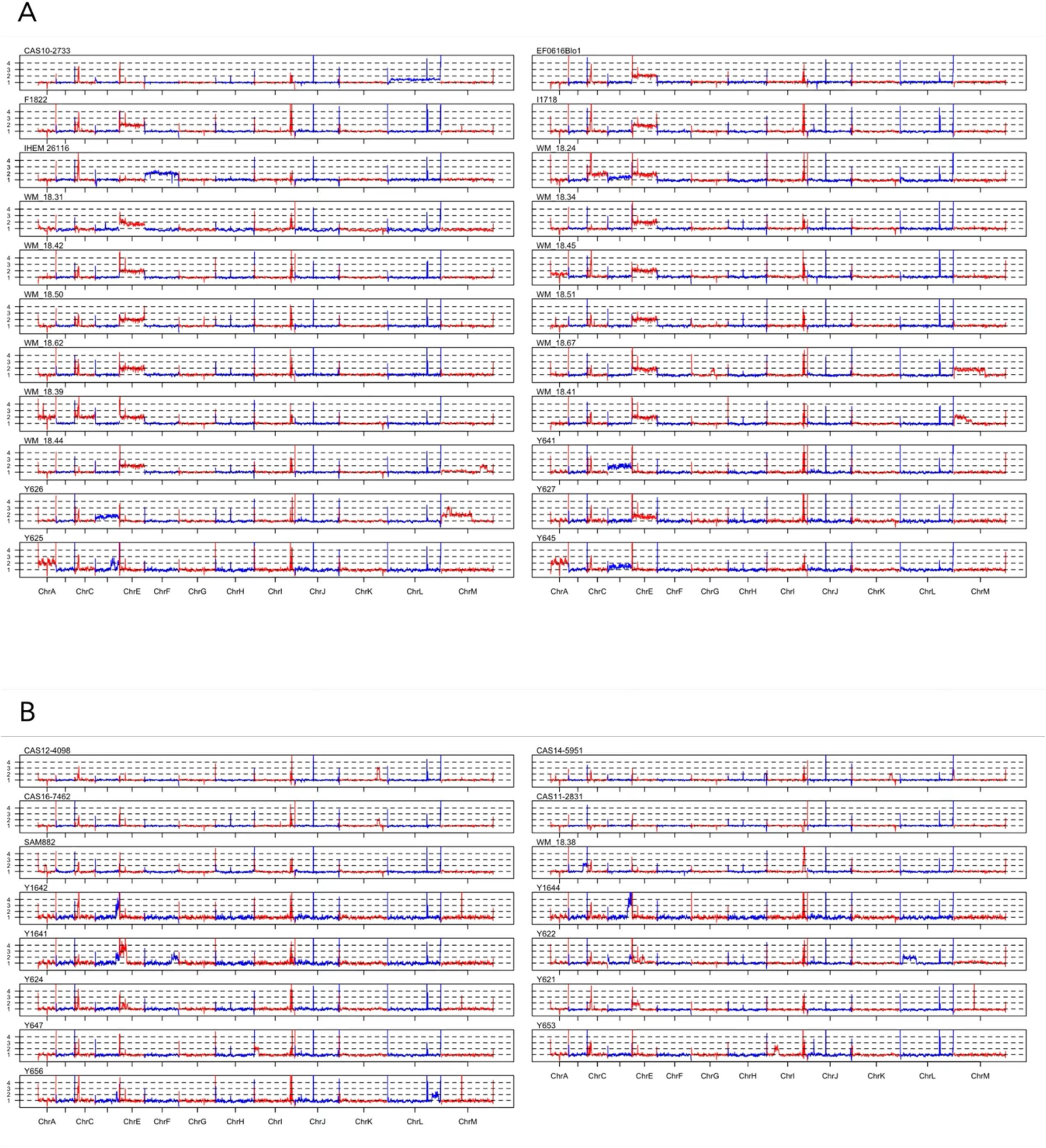
Whole-chromosome karyotypic variation among *N. glabratus* isolates. Coverage was estimated using BEDTools genomecov, and average read depth was calculated over 5 kb sliding windows per chromosome using a custom R script. Each panel shows normalized coverage per chromosome for an individual isolate, highlighting variation in chromosome copy number. Chromosomes A–M are shown from left to right, and dashed horizontal lines indicate ploidy levels (e.g., 1×, 2×, 3×). (A) Aneuploid isolates, showing whole-chromosome copy number variation. (B) Isolates with segmental copy number variation (CNV). Note that some aneuploid isolates also exhibit CNV.

**Figure S4.**
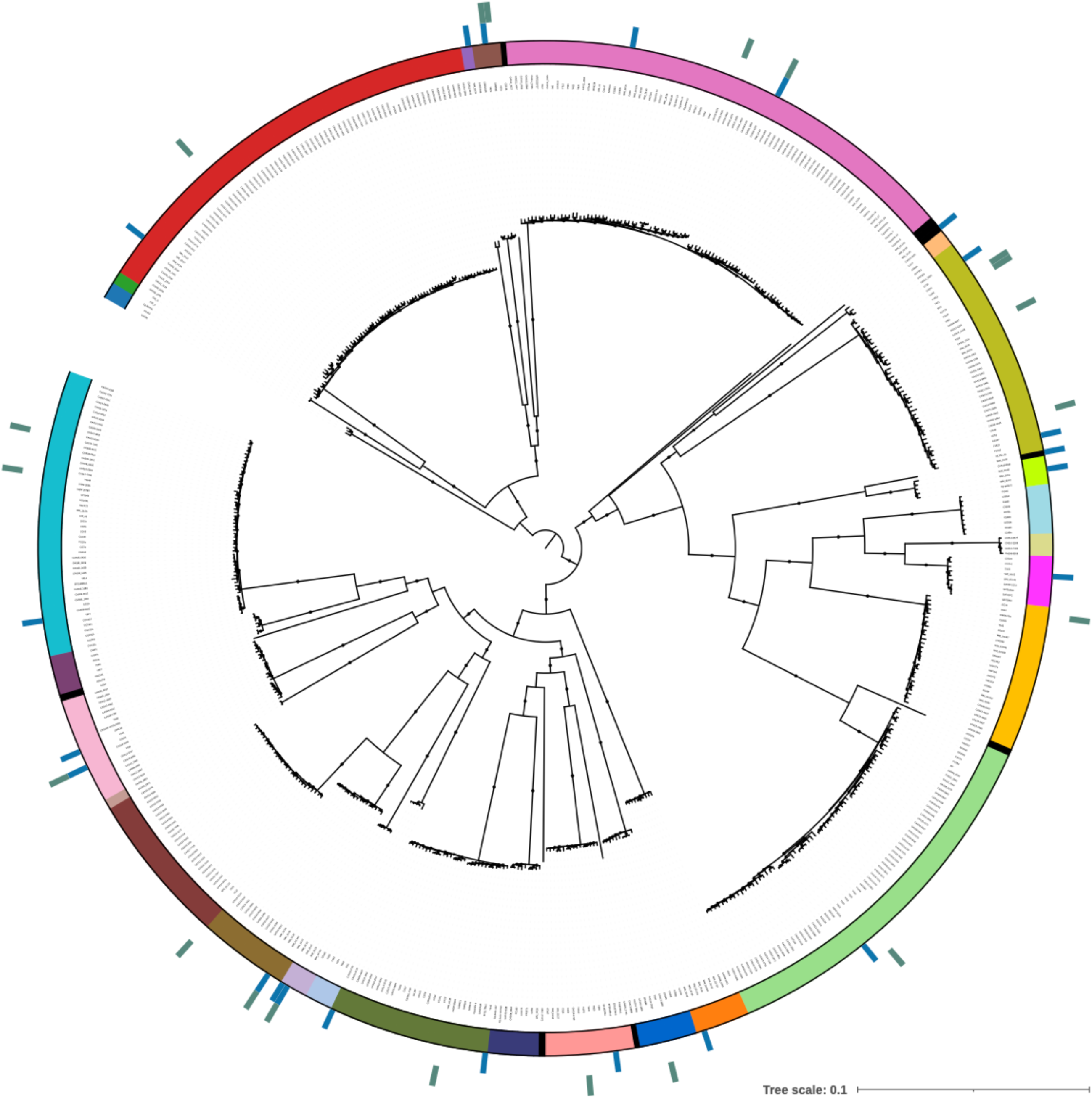
Distribution of isolates exhibiting karyotypic variation. The inner ring is colored by cluster assignment. The middle ring denotes isolates with aneuploidies, while the outer ring indicates isolates with regional copy number variations.

**Figure S5.**
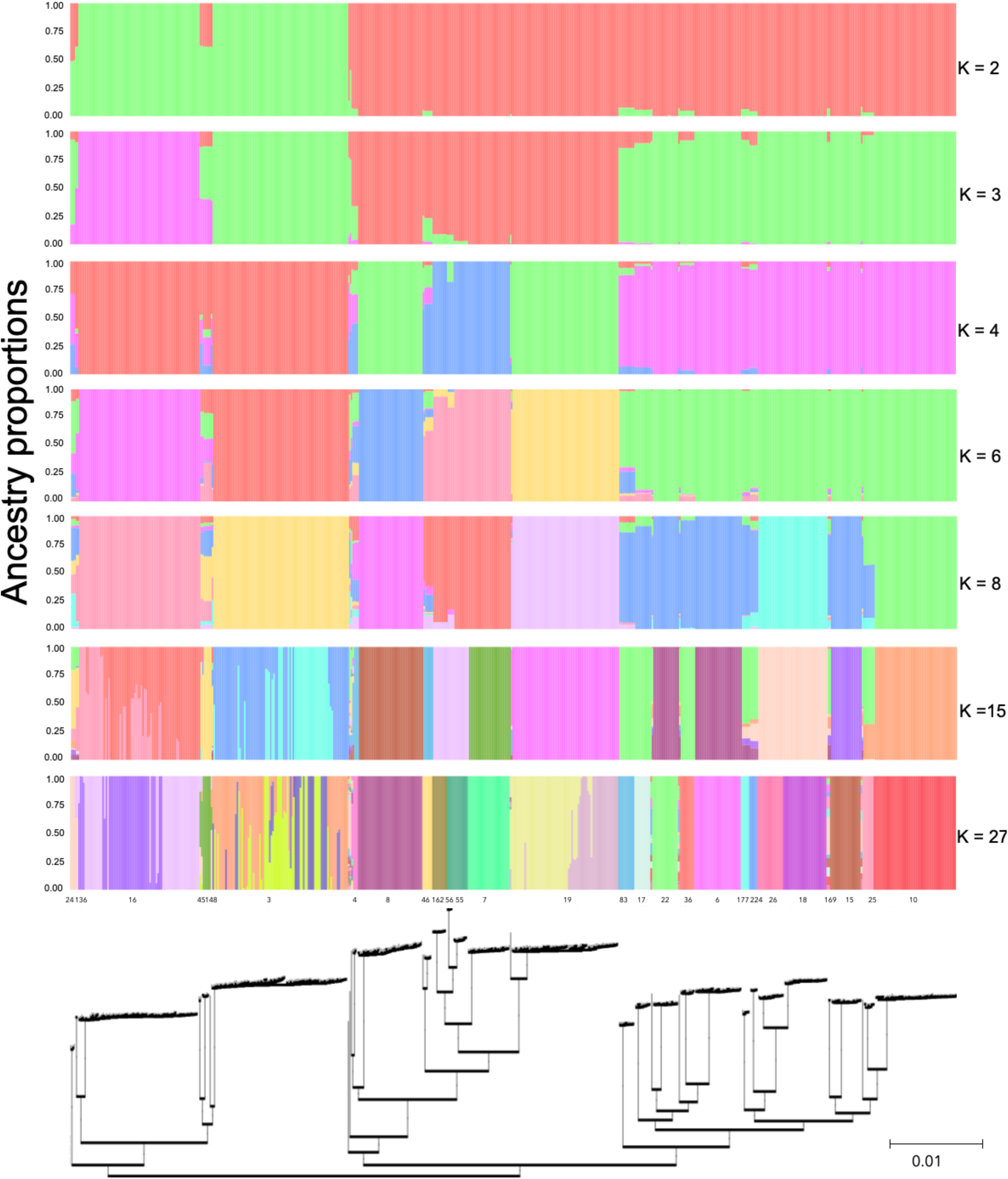
Admixture plots for all K-values (K = 2, 5, 10 16, and 27) with the corresponding phylogeny showing the positions of the isolates.

